# Untargeted metabolome- and transcriptome-wide association study identifies causal genes modulating metabolite concentrations in urine

**DOI:** 10.1101/2020.05.22.110197

**Authors:** Reyhan Sönmez Flitman, Bita Khalili, Zoltan Kutalik, Rico Rueedi, Sven Bergmann

## Abstract

In this study we investigate the results of a metabolome- and transcriptome-wide association study to identify genes influencing the human metabolome. We used RNAseq data from lymphoblastoid cell lines (LCLs) derived from 555 Caucasian individuals to characterize their transcriptome. As for the metabolome we took an untargeted approach using binned features from ^1^H nuclear magnetic resonance spectroscopy (NMR) of urine samples from the same subjects allowing for data-driven discovery of associated compounds (rather than working with a limited set of quantified metabolites).

Using pairwise linear regression we identified 21 study-wide significant associations between metabolome features and gene expression levels. We observed the most significant association between the gene *ALMS1* and two adjacent metabolome features at 2.0325 and 2.0375 ppm. By using our previously developed metabomatching methodology, we found N-Acetylaspartate (NAA) as the potential underlying metabolite whose urine concentration is correlated with *ALMS1* expression. Indeed, a number of metabolome- and genome-wide association studies (mGWAS) had already suggested the locus of this gene to be involved in regulation of N-acetylated compounds, yet were not able to identify unambiguously the exact metabolite, nor to disambiguate between *ALMS1* and *NAT8*, another gene found in the same locus as the mediator gene. The second highest significant association was observed between *HPS1* and two metabolome features at 2.8575 and 2.8725 ppm. Metabomatching of the association profile of *HPS1* with all metabolite features pointed at trimethylamine (TMA) as the most likely underlying metabolite. mGWAS had previously implicated a locus containing *HPS1* to be associated with TMA concentrations in urine but could not disambiguate this association signal from *PYROXD2*, a gene in the same locus. We used Mendelian randomization to show for both *ALMS1* and *HPS1* that their expression is causally linked to the respective metabolite concentrations.

Our study provides evidence that the integration of metabolomics with gene expression data can support mQTL analysis, helping to identify the most likely gene involved in the modulation of the metabolite concentration.

## Introduction

Genome-wide association studies (GWAS) have identified thousands of common variants that are associated with complex traits [1], but the regulatory mechanisms behind these associations mostly remain poorly understood. Pinpointing causal variants is difficult, since the lead variants associated with a trait are often in high linkage disequilibrium (LD) with other variants in the same region with only a slightly lower association signal. Such associated LD blocks typically contain several genes or functional elements, preventing the accurate identification of causal genes. Furthermore, some trait associated variants fall into intergenic regions of the genome with no obvious functional role at all.

A number of studies reported that trait associated genetic variants are significantly enriched in expression quantitative trait loci (eQTLs), suggesting that many trait associated variants affect the phenotype by altering gene expression [2–5]. There is also a growing body of literature highlighting the more pronounced effects of genetic variants on molecular traits compared to phenotypic traits [6–9]. This is not surprising, since molecular traits representing fundamental biological processes such as gene expression and metabolism are intermediates in the genotype to trait causality chain.

With high-throughput measurements becoming more accessible and widespread, integration of molecular traits into association studies has become a central challenge in the field. Such synthesis allows investigating the interplay between different organisational layers of a biological system. Despite metabolism and gene expression regulation both being fundamental biological processes that are commonly studied as molecular phenotypes, there are very few studies in humans that focus on the interplay between them. Several studies investigated the relationship between untargeted serum metabolites and whole blood gene expression in humans [10–12], but, to the best of our knowledge no transcriptome- and metabolome-wide association study has been performed using urine metabolome data of healthy human subjects.

Most metabolome- and genome-wide association studies (mGWAS) reporting metabolite quantitative trait loci (mQTL) use targeted approaches where the concentrations of a limited number of metabolites are estimated from the metabolome data generated by mass spectrometry or NMR spectroscopy. This targeted approach is limited to the number of known quantifiable metabolites in the biofluid under study. In the current study we adopted an untargeted approach, making use of the entire metabolomic data captured by binned ^1^H NMR spectra as our molecular traits. So here we present an untargeted metabolome- and transcriptome-wide association study using the entire NMR spectral information to characterize the urine metabolomes of 555 subjects and RNAseq data of lymphoblastoid cell lines (LCLs) derived from the same set of individuals. LCL have been widely used in genomic studies and proven their worth as faithful surrogates of primary tissues for studying both gene expression variation among individuals and the genetic architecture underlying regulatory variation of gene expression [13–16]. LCLs thus present an interesting system whose genetic variance in expression resembles that of the cell types affecting the urine metabolome, with the added advantage of not being influenced by immediate environmental factors such as recent changes in the diet or exposure to a drug. Despite having limited statistical power and using surrogate tissue, we identified two strong associations between gene expression levels and urine metabolome features, which allowed us to refine previous links between the corresponding genes and metabolites.

## Materials and Methods

### Study samples

Our 555 transcriptomics and metabolome profiles were measured in a randomly selected subset of individuals from CoLaus (Cohorte Lausannoise), a population-based cross-sectional study of 6,188 participants residing in Lausanne, Switzerland [17]. Recruitment to the cohort was done on the basis of a simple, non-stratified random selection of the entire Lausanne population aged 35 to 75 in 2003. The 555 samples selected for this study had a mean age of 55 (min=35, max=75) and 53% of them were women.

### Metabolomics data

We used two metabolomics data sets; the first dataset was acquired at baseline for 555 subjects and the second dataset was acquired five years later for a subset of 301 subjects. Baseline urinary metabolic profiles were generated using one-dimensional proton nuclear magnetic resonance (NMR) spectroscopy. NMR spectra were acquired at 300 K on a Bruker 16.4 T Avance II 700 MHz NMR spectrometer (Bruker Biospin, Rheinstetten, Germany) using a standard ^1^H detection pulse sequence with water suppression. The spectra were referenced to the TSP signal and phase and baseline corrected. We binned the spectra into chemical shift increments of 0.005 ppm, obtaining metabolome profiles of 2,200 metabolome features, of which 1,276 remain after filtering for missing values [18]. Lastly, the dataset was log10-transformed and standardised first across features and then samples, to make samples and feature intensities comparable.

The follow-up data was acquired with an Avance III HD 600 NMR spectrometer. These spectra were referenced to the TSP signal and phase and baseline corrected. We binned the chemical shifts into 0.005 ppm bins. After removing water and urea spectral regions (4.55-5.00 ppm and 5.5-6.1 ppm), the dataset was log10-transformed and standardised first across features then samples, to make samples and feature intensities comparable. Lastly, we performed principal component analysis (PCA) to detect outliers and 33 spectra with components scores below/above 3.5 standard deviations from the average of all components scores were removed. Our final metabolic dataset includes 1,289 features.

### Gene expression data

Total RNA was extracted from Epstein–Barr-virus-transformed lymphoblastoid cell lines (LCLs) by following the Illumina TruSeq v2 RNA Sample Preparation protocol (Illumina, Inc., San Diego, CA) by the Department of Genetic Medicine and Development at the University of Geneva. Next mRNA sequencing was performed on the Illumina HiSeq2000 platform producing 49bp paired-end reads. Paired-end reads were mapped to human genome assembly GRCh37 (hg19) with GEMTools using GENCODE v15 as gene annotation [19]. The reads were then filtered for correct orientation of the two ends and a minimum quality score of 150 while allowing 5 mismatches at both ends. Gene level read counts were quantified with an in-house script. This resulted in expression profiles of 45,470 genes for 555 individuals, which were quantified as RPKM values. Later, we transformed RPKM values by applying log-transformation [log_2_(1+RPKM)] and then standardisation across samples to make genes comparable. For our analysis we removed all genes on the sex chromosomes, as well as mitochondrial DNA genes from the gene expression data, resulting in 43,614 genes to use in the association analysis.

### Genotypic data

Genotyping was performed by using the Affymetrix GeneChip Human Mapping 500 K array set and the imputation was carried out for HapMap II SNPs. Further details of genotype calling and the imputation can be found in [18].

### Association analysis

All statistical analyses were performed using Matlab [20]. Urine metabolome features were rank normalized in order to have comparable intensities before they were used as response variables in regression.

We used a linear regression model for each pair of metabolome feature (as the response variable) and gene expression level (as the explanatory variable). The model also included the following common confounding factors: age, sex, the first four principal components of the genotypic data (correcting for population stratification) and the first 10 principal components of the gene expression data (correcting for potential batch effects). We tested 1,276 metabolome features for association with the expression of 19,123 protein coding and 24,491 non-coding genes. For the completeness of the analysis we did not apply any a-priori exclusion criteria to remove genes from the analysis. As a consequence, the distribution of genes RPKM values with significant associations were evaluated to ensure close to normality distribution for accurate regression estimations. We applied a nominal Bonferroni threshold for multiple testing p_max_ = 0.05/(125×1,109) = 3.6 ×10^−7^ by taking into account the effective number of tests which we estimated to be 125 for metabolome features and 1,109 for genes (i.e. the number of principal components explaining more than 95% of the data [21]). Only associations with p-value below p_max_ were considered significant.

### Metabomatching

Metabomatching is a method to identify metabolites underlying associations of SNPs with metabolome features [18, 22]. It compares the association profile of a given variable with all metabolome features across the full ppm range, so-called pseudospectrum, with NMR spectra of pure metabolites available in public databases such as HMDB [23]. For each metabolite m, metabomatching defines a set of features *F*_δ_(*m*) that contains all the features f that fall within a *δ* ppm vicinity of any NMR spectrum peak of m listed in the database. Metabomatching then computes the sum

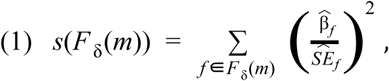

where 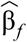 and 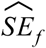 are the point estimates of feature *f* effect size and its standard error. Assuming a *χ*^2^ -distribution for the sum with |*F*_δ_(*m*)| degrees of freedom, metabomatching defines a score for each *m* as the negative logarithm of the nominal p-value corresponding to the observed sum. These scores are calculated for all the metabolites with ^1^H NMR spectrum in the database, allowing to rank them based on their likelihood to underlie the association of the variable with the metabolome features.

Although metabomatching was originally developed to use SNP-metabolome associations, recently it has been shown that it can also use co-varying features of metabolome data itself to identify metabolites [24]. In the present study we use metabomatching to identify metabolites that are associated with gene expression.

### Mendelian randomization

We performed Mendelian randomization (MR) analysis [25, 26] to assess the causal relationship between gene expression and metabolite concentration. While we used SNPs as instrumental variables (IVs), gene expression and metabolome features were interchangeably used as exposure and outcome to determine the direction of causality. For the MR analysis, we used summary statistics from mQTL/eQTL studies with higher statistical power [27, 28]. Causal effects were estimated by using the Wald method where the effect of a genetic variant on the outcome is divided by the effect of the same genetic variant on the exposure [29]. Next, ratio estimates from different instruments (SNPs) were combined by the inverse variance weighted method (IVW) to calculate the causal estimate [30].

We selected significant SNPs from relevant eQTL/mQTL studies as our IVs. To detect the independent SNPs, we used a stepwise pruning approach where we first selected the strongest lead eQTL/mQTL and then pruned the rest of the SNPs in a stepwise manner if they were correlated with the lead SNP (r^2^ > 0.2). We repeated the pruning process with the next available SNP until there were no SNPs left to prune. We used Cochran’s Q test to determine heterogeneity among the candidate instruments [31]. The SNPs were pruned in a stepwise manner from the model until the model did not show any more signs of heterogeneity (Cochran’s Q statistic p-value > 0.05/#of original instruments). We also applied more robust MR analysis methods than IVW, such as the median estimator and MR-Egger regression to evaluate the significance of the causal estimates [32]. These methods are known to have more relaxed MR assumptions and they can tolerate the violation of the exclusion-restriction assumption for some instruments. For all MR analysis we used the Mendelian Randomization package implemented in R [33].

## Analysis & Results

### Association analysis

We performed an untargeted metabolome- and transcriptome-wide association study by pairwise linear regression of log-transformed expression levels of each of the 43,614 genes (as response variable) onto each of the 1,276 metabolome features (as explanatory variable). The metabolome features resulted from binning the raw urinary NMR spectra with a bin-size of 0.005 ppm, and rank normalizing each bin passing QC (see Methods). The gene expression levels, quantified as RPKM, were measured using RNAseq on lymphoblastoid cell lines derived from the same set of 555 subjects.

Figure 1 shows the qq-plot of all pairwise associations. It is well calibrated, and only two association p-values (both involving the *ALMS1* gene, see below) are highly significant (FDR<0.05). Yet, applying an adjusted Bonferroni threshold of 3.6 ×10 ^−7^ to account for the effective number of independent variables (see Methods) we identified 25 additional marginally significant feature-gene associations. The 27 association pairs involved 22 unique genes and 25 unique features. We did not apply any a-priori exclusion criteria to remove genes from the analysis. Instead, we inspected the expression value distributions of these 22 significant genes in order to identify cases in which the small p-value may be due to a problematic distribution of the expression values. Indeed, we observed that some of the genes had zero expression values for a sizable fraction of the samples and very low expression values otherwise. Based on the distributions we filtered out all genes that had more than 95% RPKM values equal to 0 and a maximum RPKM value over all samples lower than 1. Applying this rather mild filtering removed 11,547 out of the 43,614 autosomal genes (26%) and 1,994 out of 19,123 protein-coding genes (10%). Amongst the 22 marginally significant associations five (23%) were removed. Expression distributions of the discarded as well as remaining genes are presented in Supplementary Figure 1 and 2, respectively. We report the remaining 21 significant associations corresponding to 17 unique genes and 19 unique features in Table 1.

**Table 1:**
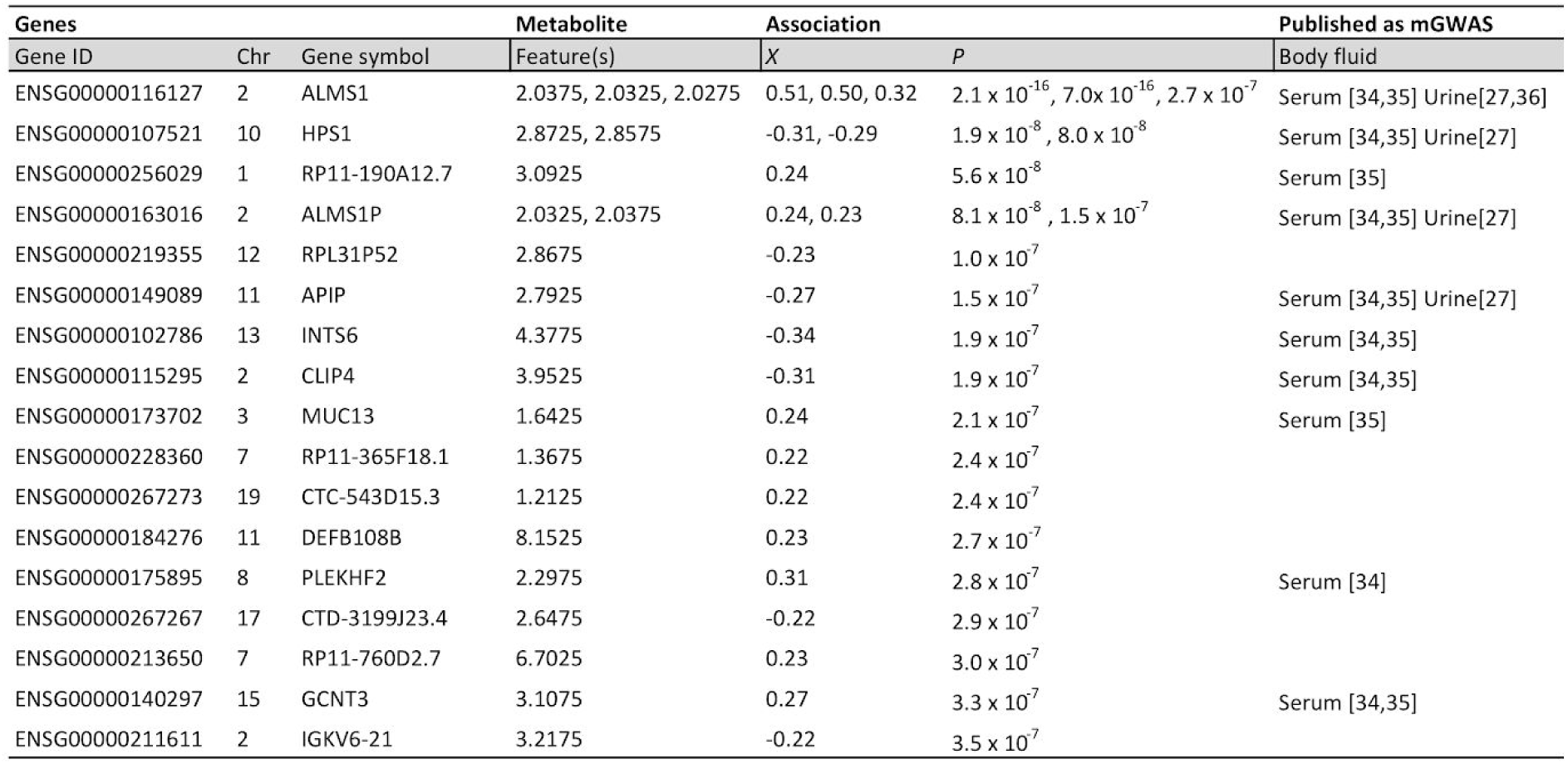
21 study-wide significant associations from metabolome- and transcriptome-wide association analysis, corresponding to 17 unique genes and 19 unique features. Abbreviations: GeneID – Ensembl Gene ID (NCBI build 37), Chr – chromosome, *X* – effect size, *P* – P-value.

| Genes |  |  | Metabolite | Association |  | Published as mGWAS |
| --- | --- | --- | --- | --- | --- | --- |
| Gene ID | Chr | Gene symbol | Feature(s) | X | P | Body fluid |
| ENSG00000116127 | 2 | ALMS1 | 2.0375, 2.0325, 2.0275 | 0.51, 0.50, 0.32 | $2.1 \times 10^{-16}$, $7.0 \times 10^{-16}$, $2.7 \times 10^{-7}$ | Serum [34,35] Urine[27,36] |
| ENSG00000107521 | 10 | HPS1 | 2.8725, 2.8575 | -0.31, -0.29 | $1.9 \times 10^{-8}$, $8.0 \times 10^{-8}$ | Serum [34,35] Urine[27] |
| ENSG00000256029 | 1 | RP11-190A12.7 | 3.0925 | 0.24 | $5.6 \times 10^{-8}$ | Serum [35] |
| ENSG00000163016 | 2 | ALMS1P | 2.0325, 2.0375 | 0.24, 0.23 | $8.1 \times 10^{-8}$, $1.5 \times 10^{-7}$ | Serum [34,35] Urine[27] |
| ENSG00000219355 | 12 | RPL31P52 | 2.8675 | -0.23 | $1.0 \times 10^{-7}$ | |
| ENSG00000149089 | 11 | APIP | 2.7925 | -0.27 | $1.5 \times 10^{-7}$ | Serum [34,35] Urine[27] |
| ENSG00000102786 | 13 | INTS6 | 4.3775 | -0.34 | $1.9 \times 10^{-7}$ | Serum [34,35] |
| ENSG00000115295 | 2 | CLIP4 | 3.9525 | -0.31 | $1.9 \times 10^{-7}$ | Serum [34,35] |
| ENSG00000173702 | 3 | MUC13 | 1.6425 | 0.24 | $2.1 \times 10^{-7}$ | Serum [35] |
| ENSG00000228360 | 7 | RP11-365F18.1 | 1.3675 | 0.22 | $2.4 \times 10^{-7}$ | |
| ENSG00000267273 | 19 | CTC-543D15.3 | 1.2125 | 0.22 | $2.4 \times 10^{-7}$ | |
| ENSG00000184276 | 11 | DEFB108B | 8.1525 | 0.23 | $2.7 \times 10^{-7}$ | |
| ENSG00000175895 | 8 | PLEKHF2 | 2.2975 | 0.31 | $2.8 \times 10^{-7}$ | Serum [34] |
| ENSG00000267267 | 17 | CTD-3199J23.4 | 2.6475 | -0.22 | $2.9 \times 10^{-7}$ | |
| ENSG00000213650 | 7 | RP11-760D2.7 | 6.7025 | 0.23 | $3.0 \times 10^{-7}$ | |
| ENSG00000140297 | 15 | GCNT3 | 3.1075 | 0.27 | $3.3 \times 10^{-7}$ | Serum [34,35] |
| ENSG00000211611 | 2 | IGKV6-21 | 3.2175 | -0.22 | $3.5 \times 10^{-7}$ | |

**Figure 1:**
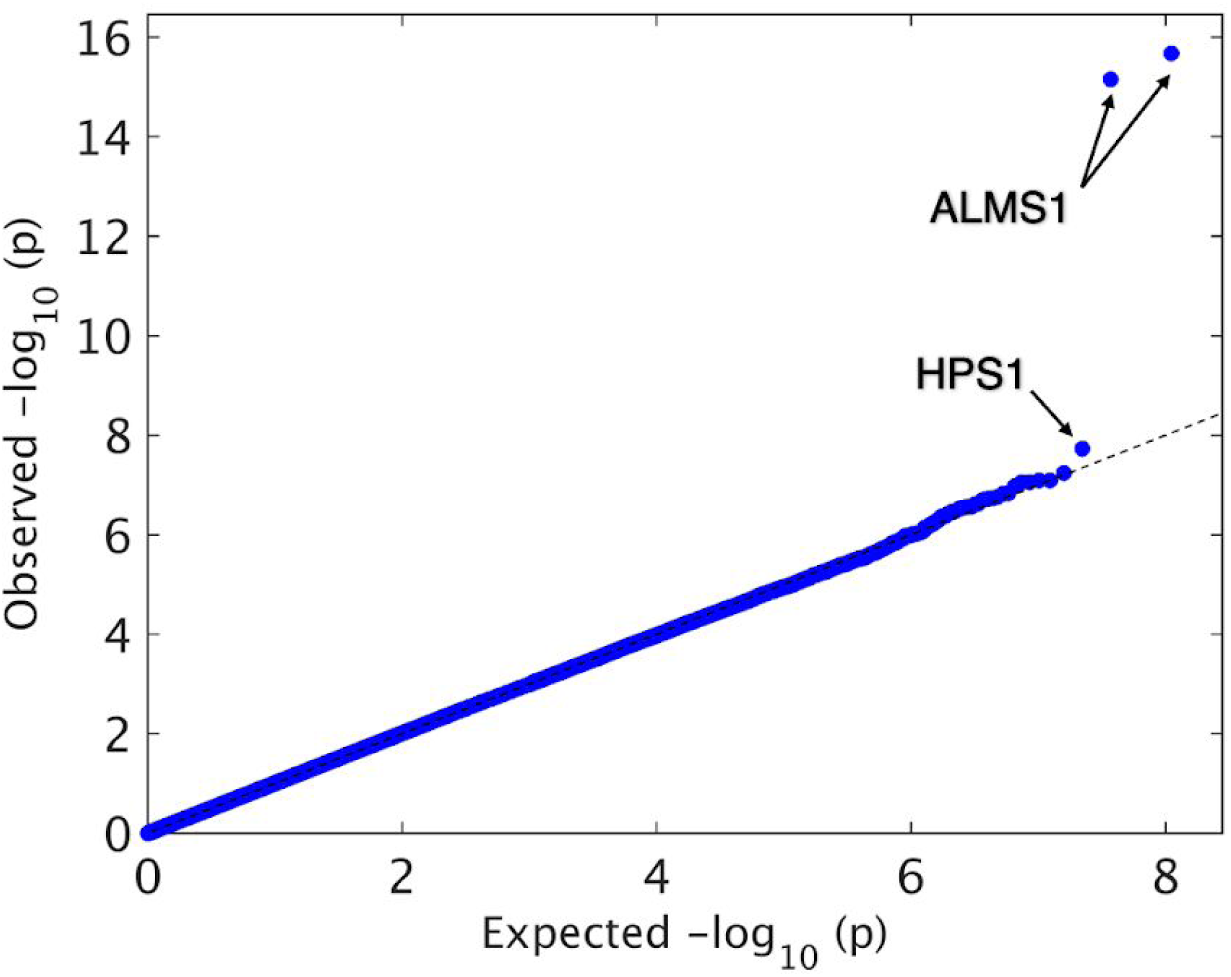
QQ-plot showing −log10(p)-values of metabolome- and transcriptome-wide association analysis. The features that significantly associate with *ALMS1* expression are ranking as 1st, 2nd and 8th; the features associated with *HPS1* expression are ranking as 3rd and 5th and the features associated with *ALMS1P* expression are ranking as 6th and 7th.

### Metabolite discovery

To find the metabolites underlying these significant associations between gene expression levels and metabolome features we used metabomatching. Metabomatching has been previously established as an effective tool for prioritizing candidate metabolites underlying SNP-metabolome features association profiles, so-called pseudospectra [18, 27]. In this study we used association profiles of genes which had at least one significantly associated metabolite feature as input to metabomatching and found that the pseudospectra of *ALMS1* and *ALMS1P* matched well with the N-Acetylaspartate (NAA) NMR spectrum and that the pseudospectrum of *HPS1* matched well with the trimethylamine (TMA) NMR spectrum (Figure 2).

**Figure 2:**
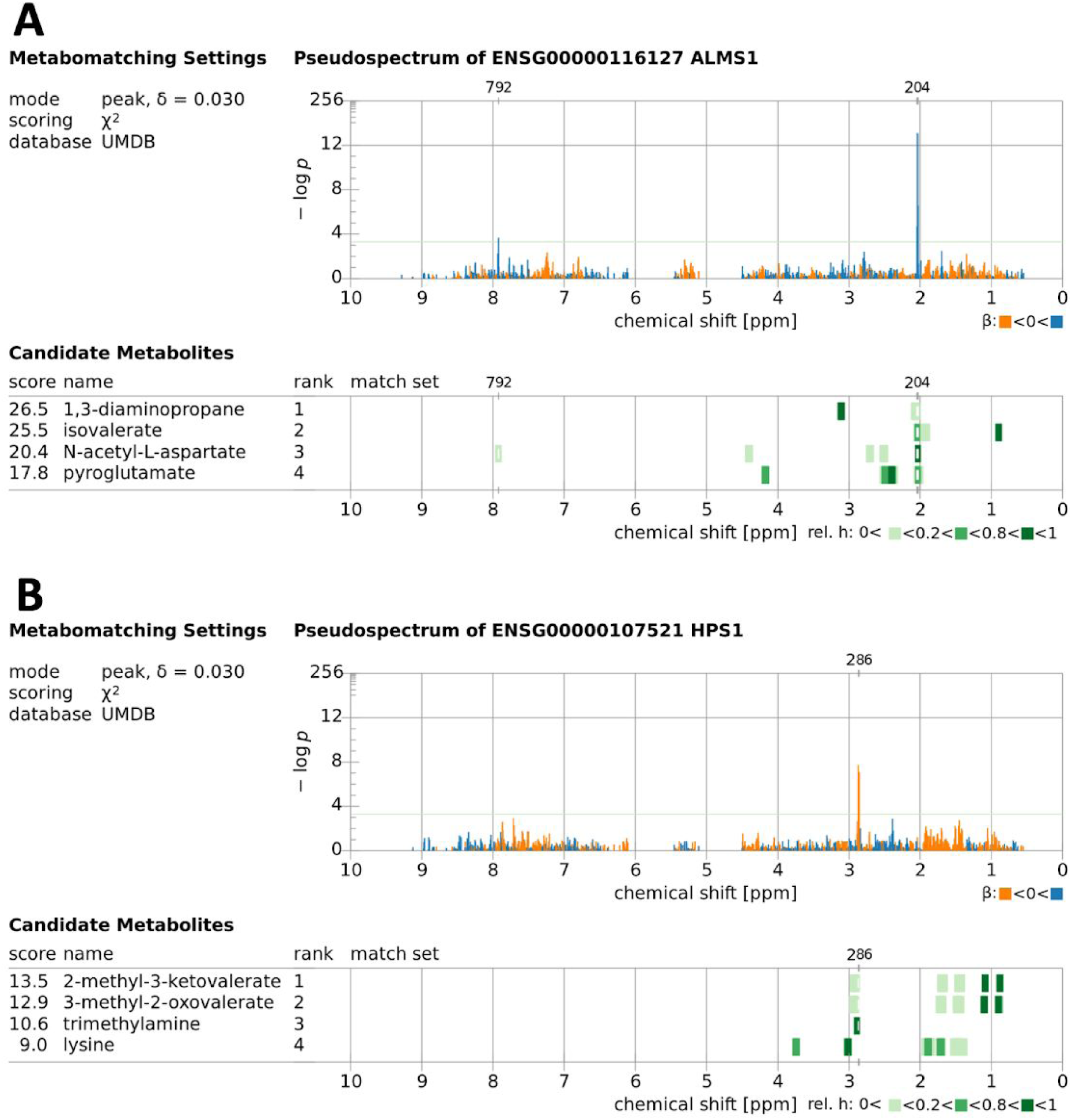
Metabomatching figures showing the pseudospectra derived from gene expression – metabolome features associations [22]. The features in each pseudospectrum are color-coded by the sign of the effect size and the four highest ranking candidate metabolites are listed on the lower left with their reference NMR spectra shown on the right (color coding indicating their relative peak intensities). A) CoLaus urine metabolome-*ALMS1* gene expression association profile metabomatching figure. Leading features allowing metabolite identification are at 2.03 ppm and 7.92 ppm regions which match well with the highest intensity peak of NAA and one of the lower intensity peaks of the NAA NMR spectrum, respectively. B) CoLaus urine metabolome – *HPS1* gene expression association profile metabomatching figure. Leading features allowing metabolite identification are at 2.87 and 2.86 ppm which match well with TMA singlet.

As shown in Table 1, the expression of *ALMS1* significantly associates with three neighboring features at 2.0375 ppm (p-value= 2×10 ^−16^), 2.0325 ppm (p-value=7 ×10 ^−16^) and 2.0275 ppm (p-value=3 ×10 ^−7^). There are few metabolites with resonances in this region and usually a singlet signal in this area is interpreted as the N-acetylated resonance detected in the ^1^H NMR spectrum of N-acetylated compounds [37]. As illustrated in Figure 2A, among the top three metabolites suggested by metabomatching that have a peak at 2.03 ppm, the only one with the highest intensity peak at this position is NAA. Also the presence of a secondary peak in the pseudospectrum at 7.9225 ppm matches well with one of the lower intensity peaks of the NMR spectrum of NAA reported at 7.92 ppm in HMDB, even though the association p-value of this feature is below the Bonferroni threshold (p-value=2 ×10 ^−4^). Similarly, metabomatching the pseudospectrum of *ALMS1P* (*ALMS1* pseudogene) points to NAA as the most likely matching N-acetylated compound (Supplementary Figure 3). The metabolome features pointing to NAA are the same features as in *ALMS1* but with lower association p-values (2.0375 ppm with p-value=1 ×10 ^−7^, 2.0325 ppm with p-value= 8 ×10 ^−8^, 7.9225 ppm with p-value= 2 ×10 ^−4^).

**Figure 3:**
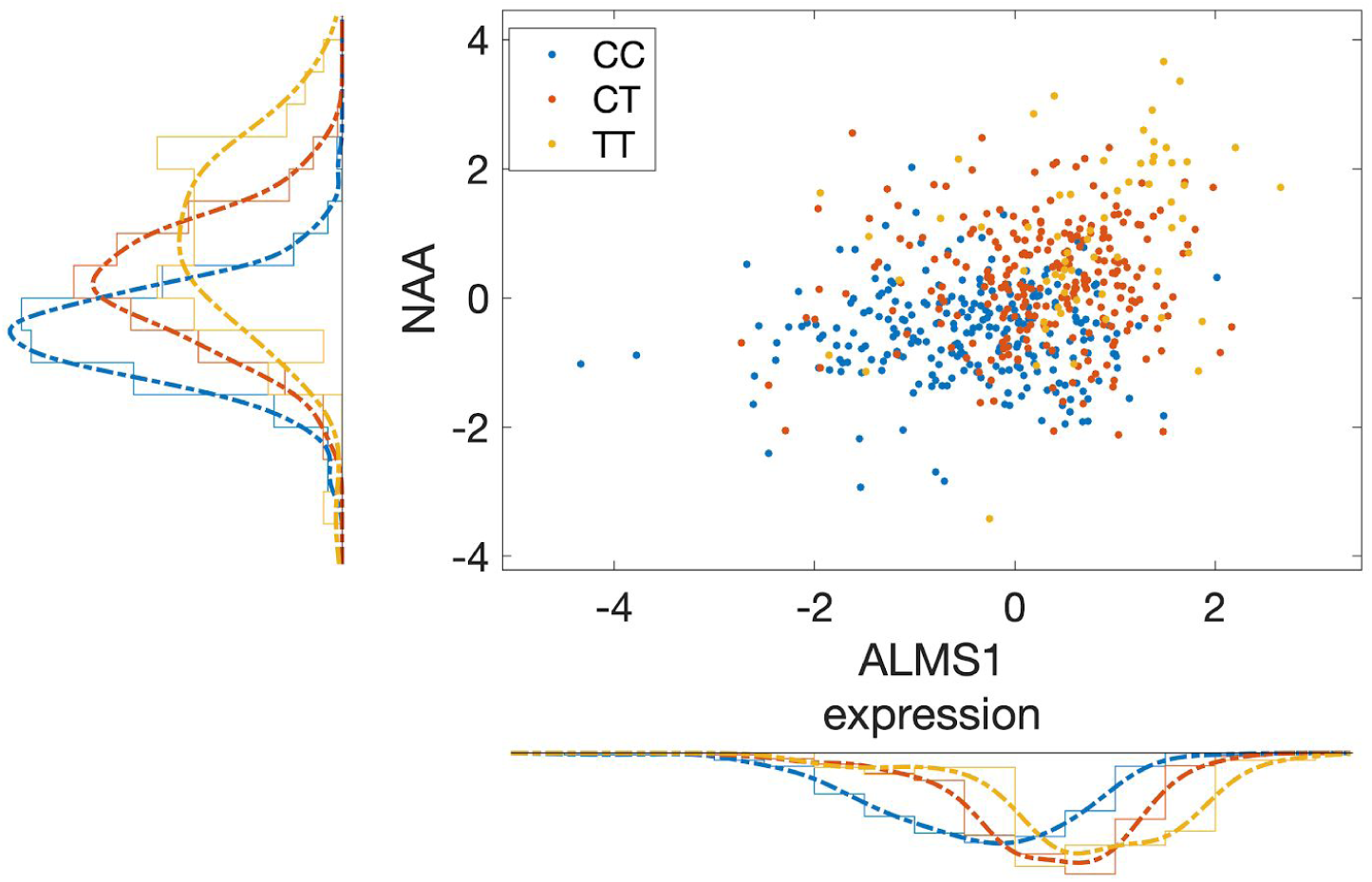
SNP rs7566315, showing a mQTL effect on NAA and an eQTL effect on *ALMS1* gene expression. Each point represents a study sample. NAA concentration is approximated by the feature at 2.0375 ppm that is log10 transformed after feature- and sample-wise z-scoring (y-axis). *ALMS1* expression is quantified as log2 transformed RPKM+1 values (x-axis). Color code represents the genotype of rs7566315 (legend).

The reference spectrum of NAA in the Urinary Metabolome Database (UMDB) that we used for metabomatching was recorded in water. In order to verify that the peaks of this spectrum are comparable to those of NAA in urine, we spiked NAA into pooled urine samples from our collection at a concentration of 10 mM and recorded its ^1^H NMR spectrum. Inspecting the 5 multiplet regions of NAA, we concluded that the NAA peak positions are very similar in both solvents (Supplementary Figure 4). To further investigate if a better match exists among all the N-acetylated family of compounds, we built a library consisting of all N-acetylated compounds proton NMR spectra available in HMDB and the Biological Magnetic Resonance Data Bank (BMRB). NAA remained the best metabomatching hit for the *ALMS1* pseudospectrum (Supplementary Figure 5). Figure 3 illustrates the relationship between *ALMS1* gene expression level and the NAA metabolite concentration where every point in the plot represents a study sample and each of the samples are color coded according to the genotype at rs7566315 SNP, that is an eQTL of *ALMS1* and mQTL of NAA.

**Figure 4:**
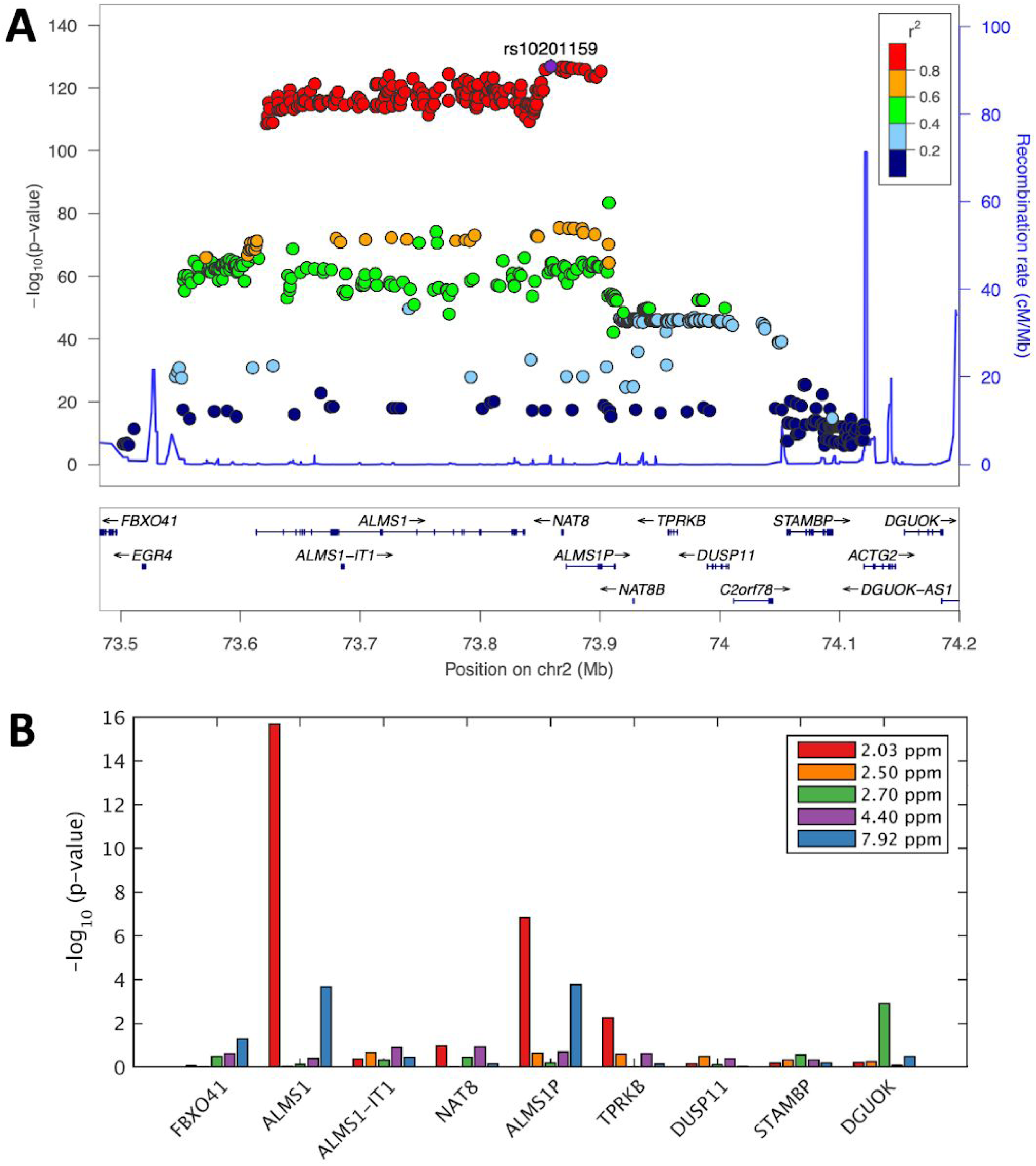
A) LocusZoom plot for *ALMS1/NAT8* locus, where the SNPs are associated with metabolome feature at 2.0375 ppm, LD colored with respect to lead mQTL. B) Bar plot shows −log_10_ transformed p-values from associating expression values of nine genes in the locus with the five NAA features.

**Figure 5:**
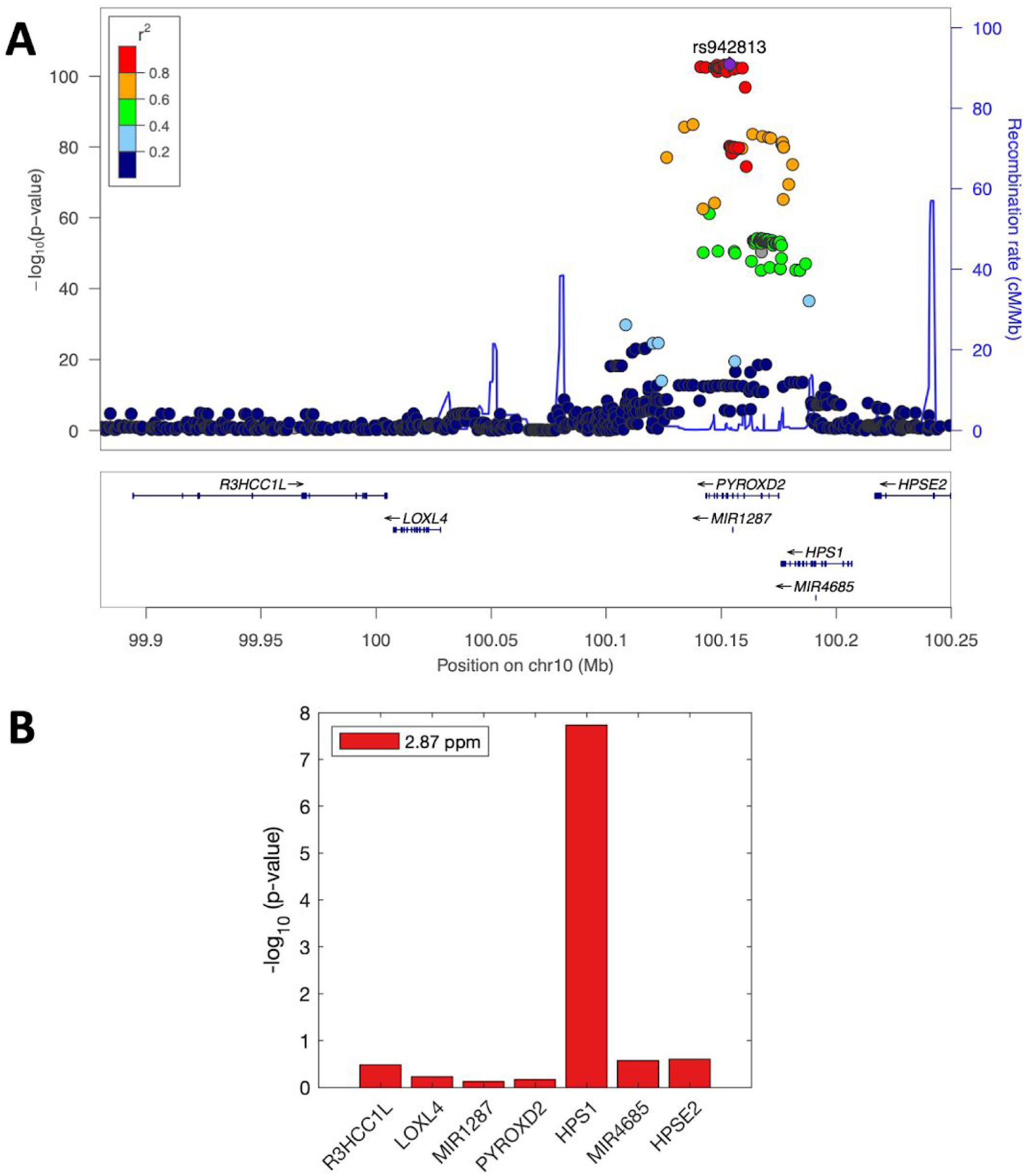
A) LocusZoom plot for *HPS1/PYROXD2* locus, where the SNPs are associated with metabolome feature at 2.8725 ppm, LD colored with respect to lead metaboliteQTL. B) Bar plot shows −log_10_ transformed p-values from associating expression values of seven genes in the locus with the TMA feature at 2.87 ppm.

The third and fifth strongest associations in Table 1 are between *HPS1* gene expression and two neighboring metabolome features at 2.8725 ppm (p-value=2 ×10 ^−8^) and 2.8575 ppm (p-value= 8 ×10 ^−8^), respectively. Figure 2B shows the metabomatching result of the *HPS1* pseudospectrum. Among the top three metabolites suggested by metabomatching, trimethylamine (TMA) is the most plausible metabolite driving the association pattern, as it is the only metabolite with its highest intensity NMR peak at 2.86 ppm region and no missing peaks. Schematic representation of the match between pseudospectra and the NMR spectra for both *ALMS1* and *HPS1* can be seen in Supplementary Figure 6.

### Validation of *ALMS1* and *HPS1* associations

To the best of our knowledge, there is no other study with urine NMR spectra and expression data of LCLs derived from the same subjects that is of comparable or larger sample size, precluding proper out-of-sample replication of our results. We have, however, access to additional urine NMR spectra from samples collected for a subset of 301 CoLaus subjects in a follow-up study conducted five years after the baseline data collection. We note that the follow-up NMR data are not independent from the baseline data, yet they were obtained from physically different samples collected at a significantly later time and processed in a different NMR spectrometer and facility. As for the expression data, we only have those from LCLs derived from blood taken at baseline, so we could only test whether the associations we observed between baseline metabolomics and baseline transcriptomics measurements would persist as associations between follow-up metabolomics and baseline transcriptomics data.

We thus asked whether our significant and marginally significant results can be confirmed also using the follow up metabolomics data. We focused on the *ALMS1* and *ALMS1P* gene expression association with NAA and the *HPS1* gene expression association with TMA. As baseline and follow-up urine NMR data were each processed and binned individually, the features did not correspond one-to-one between the studies. To test the association of these three genes with relevant features, we selected all features within +/− 0.03 ppm neighborhood of top features associated with these genes from baseline dataset; i.e. 2.0375 ppm for *ALMS1* and *ALMS1P*, and 2.8575 ppm for *HPS1*. This resulted in 12 features to test for each of the genes. We used a Bonferroni multiple testing corrected p-value threshold of 0.05/(12 features × 3 genes) = 1.4 ×10 ^−3^.

In the follow-up, *ALMS1* gene expression level significantly associated with three neighboring features at 2.042 ppm (p-value=5.1 ×10^−7^), 2.037 ppm (p-value=3.7 ×10^−6^) and 2.032 ppm (p-value=3.9 ×10^−4^), likely corresponding to the features at 2.0375 and 2.0325 ppm in the baseline association study. *HPS1* gene expression level significantly associated with 2 features at 2.869 ppm (p-value=2.2 ×10 ^−5^) and 2.859 ppm (p-value=1.3 ×10 ^−3^) that likely correspond to the features at 2.8725 and 2.8575 ppm in the baseline dataset. *ALMS1P* however did not show any significant association with candidate features in the follow-up study. Supplementary Table 1 summarises our validation results.

### Comparison with mGWAS results

We performed an association study with metabolome features in the NAA and TMA NMR peak regions using data from 826 individuals of the CoLaus cohort for whom the urinary NMR spectra are available (similar to [18]). Figure 4A shows the locuszoom figure of SNPs in loci surrounding *ALMS1*/*NAT8* locus with significant association p-values with metabolome feature at 2.0375 ppm. The SNPs most strongly associated with this metabolome feature are correlated with each other and lie within a locus containing *ALMS1*, *ALMS1-IT1*, *NAT8* and *ALMS1P* genes (r^2^>0.8). In Figure 4B, we show the p-values for association of expression values from nine genes with five different metabolome features that represent all multiplet regions of NAA (see Supplementary Figure 4 for a wider range of genes in the locus). *ALMS1* and *ALMS1P* have the most significant association results with the 2.0375 ppm feature, compared to the rest of the genes. Concordantly, *ALMS1* and *ALMS1P* gene expression levels are associated more significantly to the feature at 7.9225 ppm, the secondary feature in our NAA identification, compared to the other genes at the locus. Figure 5A shows the significant association pattern of SNPs in the loci surrounding *HPS1*/*PYROXD2* locus with metabolome feature at 2.8725 ppm and Figure 5B shows the significance level for association of expression values from seven genes with the same metabolome feature. Even though the SNPs with the most significant association with feature 2.8725 are physically located closer to *PYROXD2* gene rather than *HPS1* gene, the expression level of *PYROXD2* does not show significant association with this feature. Inspecting the list of published mGWAS in humans [38], we found that the SNPs in both *ALMS1* and *HPS1* loci have been previously reported to associate with a number of metabolic traits (Tables 2). The *ALMS1* locus has previously been associated with a number of N-acetylated compounds, while *HPS1* locus has been associated with various metabolites including trimethylamine and dimethylamine [18, 27, 36]. In mGWAS studies determining the mediator genes is not a straightforward procedure, as mQTL SNPs are indistinguishable from neighboring SNPs in LD, and mediator genes of the mQTLs are often inferred based on their physical proximity to the SNPs or functional relevance. Consequently, published mGWAS studies were not able to distinguish between *NAT8* and *ALMS1* or *HPS1* and *PYROXD2* as mediator genes of NAA and TMA, respectively. In contrast, in the current association study we use gene expression data allowing us to pinpoint *ALMS1* and *HPS1* as mediator genes.

**Table 2:**
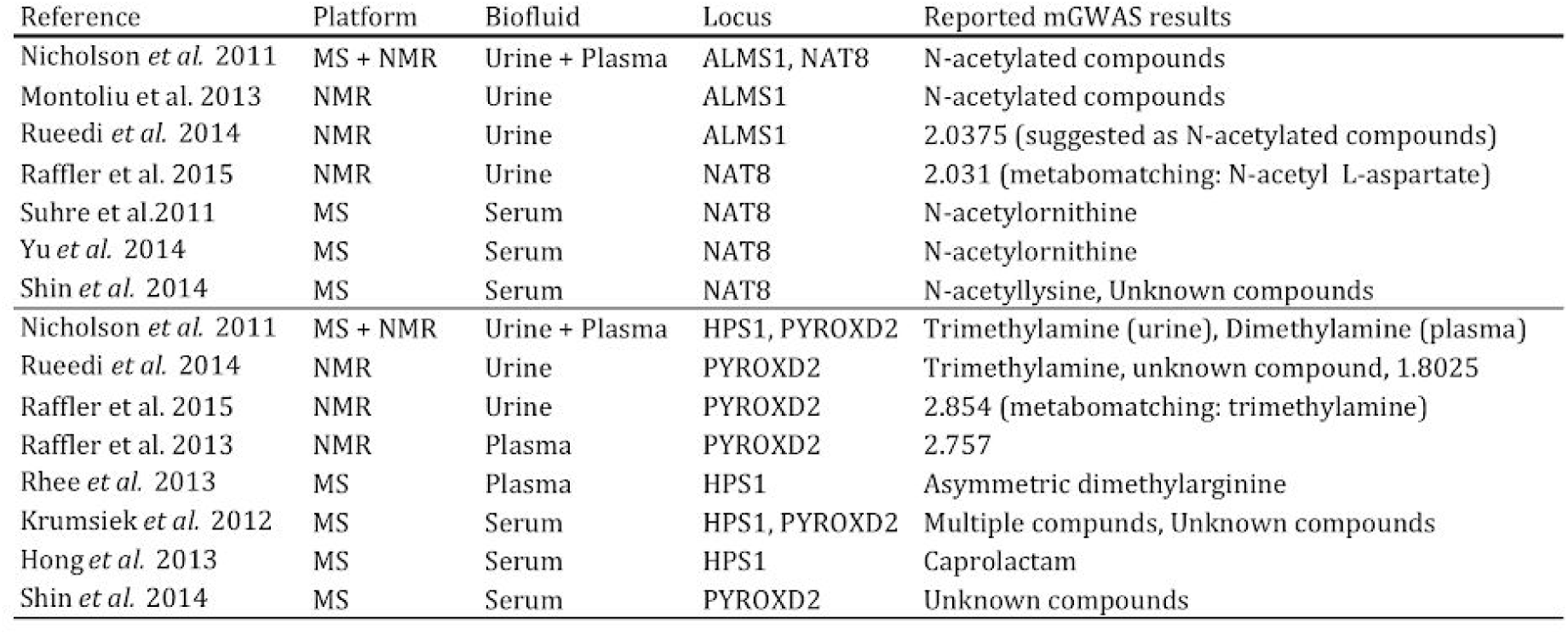
List of published mGWAS results in humans concerning *ALMS1*/*NAT8* and *HPS1*/*PYROXD2* loci. MS:Mass Spectrometry, numbers in reported mGWAS results section refer to NMR spectral shift positions in ppm.

To further evaluate the possible regulation of NAA and TMA by other genes suggested by published mGWAS studies, we investigated the metabomatching plots of these genes in order to see if they pointed to any N-acetylated compounds/TMA. The investigated genes either (a) were the target of an eQTL SNP that is mQTL of NAA/TMA, or (b) were within 500kb of *ALMS1*/*HPS1*. However none of these candidate genes produced a pseudospectrum containing even a single nominally significant signal pointing to NAA/TMA.

### Causality analysis

We performed MR analysis using summary statistics from the eQTLGen Consortium [28] and Raffler et al. [27] for eQTL and untargeted mQTL results, respectively. We investigated both the causal effect of the gene expression on the metabolite concentration and vice versa for the *ALMS1*-NAA and *HPS1*-TMA gene-metabolite pairs.

In the MR analysis where we investigated the causal effect of *ALMS1* gene on NAA concentration, instrumental variables (IVs) were selected among the SNPs that were reported as significant eQTLs (FDR<0.05) in eQTLGen and that were also measured in Raffler et al., resulting in 86 SNPs. By applying the stepwise pruning approach (see Methods) we found 14 independent SNPs as candidate IVs. Next, we performed Cochran’s Q test to detect heterogeneity among these 14 SNPs and removed a further three of those, resulting in 11 SNPs as potentially valid IVs to use in the MR analysis (see Methods). As for the outcome, we used NMR peak intensities as proxies for the concentration of NAA as there were no targeted studies reporting summary statistics explicitly for NAA concentration. To this end we used the peak at 2.0308 ppm reported in Raffler et al., as this peak is the highest peak in the NAA spectrum and often used to estimate the concentration of N-acetylated compounds (NAC) [36, 39]. NAA has other NMR peaks in its spectrum, yet the observed intensities at these peaks are much lower and therefore difficult to detect robustly by NMR spectroscopy. Indeed these peaks were only weakly correlated amongst themselves and with the main peak at 2.03 ppm region (pearson rho<0.5), so they were too noisy to define a more robust estimate of the NAA concentration than the main peak on its own. For these reasons we decided to perform our MR analysis using only the intensity measure at 2.03 ppm as outcome, which implies therefore that we studied the causality of any NAC rather than NAA specifically. Causal effect estimates given by different meta-analysis methods are reported in Table 3. All methods agreed on *ALMS1* expression level being causal for NAC concentrations.

**Table 3:**
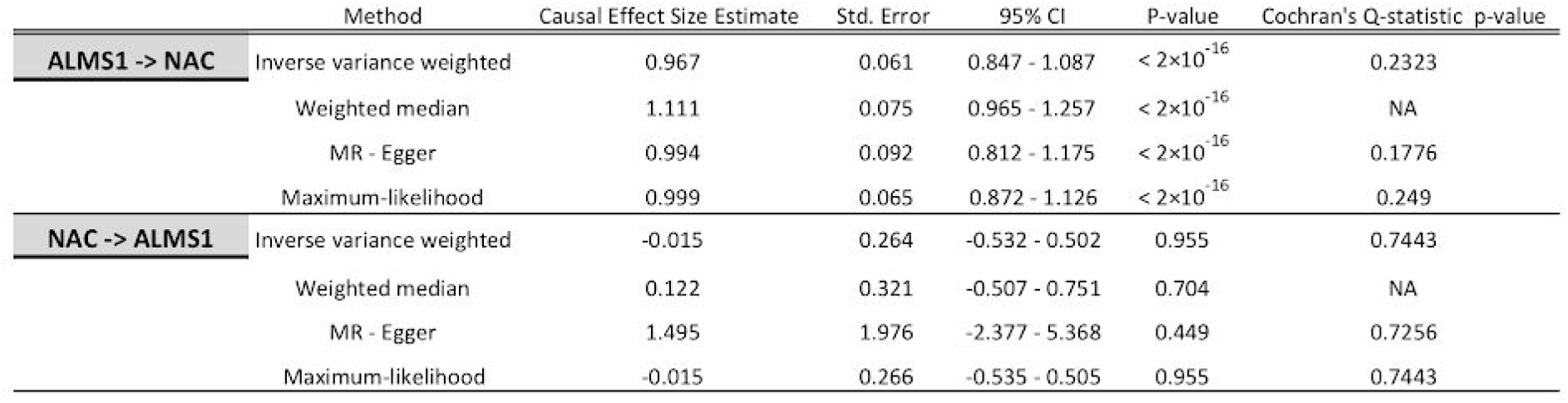
MR results for testing causal effect of *ALMS1* gene expression levels on N-acetylated compounds (*ALMS1* -> NAC) and MR results for testing causal effect of N-acetylated compounds on *ALMS1* gene expression levels (NAC -> *ALMS1*) using summary statistics data.

For the completeness of the analysis, we also tested the causal effect of NAC on *ALMS1* gene expression level. IVs were selected among the SNPs that were reported as significant mQTLs (p-value < 1 ×10 ^−6^) in Raffler et al. [27]. Amongst the cis-eQTLs of *ALMS1* from eQTLGen, most candidate IVs seemed to have direct pleiotropic effect on *ALMS1* expression in cis, reflected by the strong heterogeneity between their expected and observed effects. To overcome this problem we sought to use also trans-eQTLs of *ALMS1*, however none of the candidate IVs were measured in the trans-eQTL study of eQTLGen. As an alternative, we performed an association study between the candidate IVs and *ALMS1* gene expression level as measured in CoLaus and used these eQTL results in the MR analysis. Overall, we identified 26 significant mQTLs for the 2.03 ppm feature in Raffler et al. (p-value < 1 ×10 ^−6^) which corresponded to six independent SNPs. Two of the six candidate IVs exhibited pleiotropic effects and they were removed from the analysis. Finally, we had four SNPs as potentially valid IVs to use in the MR analysis (see Methods). Causal effect estimates given by different meta-analysis methods are reported in Table 3. None of the methods found NAC concentration to be causal for *ALMS1* gene expression level. However, it should be noted that due to low sample size of trans-eQTL study, this particular MR analysis was underpowered.

For the MR analysis of the *HPS1* gene, IVs were selected among the SNPs that were reported as significant eQTLs (FDR<0.05) in eQTLGen and that were also measured in Raffler et al. [27]. As for the outcome, similarly to NAA, there were no studies reporting targeted summary statistics for TMA concentration, therefore we used the NMR peak intensities to estimate the concentration of TMA. According to HMDB, TMA has one singlet at 2.89 ppm where the peak position ranges from 2.79 to 2.99 ppm. In the Raffler et al. dataset we used the intensity of feature at 2.8541 ppm as a proxy of TMA concentration. For the MR analysis we had 77 candidate SNPs six of which were selected as valid IVs as they were independent and did not exhibit heterogeneity (see Methods). Causal effects estimated by using different meta-analysis methods are reported in Table 4. All of the methods agreed on *HPS1* gene expression having a causal effect on TMA concentration.

**Table 4:**
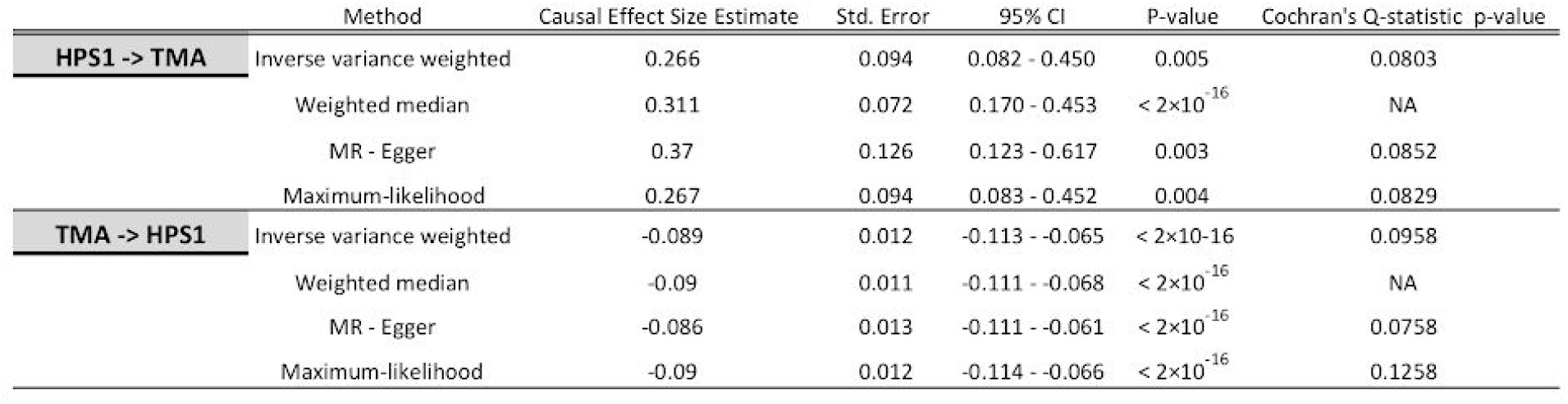
MR results for testing causal effect of *HPS1* gene expression level on TMA (*HPS1* -> TMA) and MR results for testing causal effect of TMA on *HPS1* gene expression level (TMA -> *HPS1*) using summary statistics data.

We also explored the causal effect in the other direction, testing the causal effect of TMA concentration on *HPS1* gene expression. There were 87 significant mQTLs in Raffler et al. [27] that were also measured in eQTLGen. By applying the stepwise pruning approach and removing the SNPs showing heterogeneity (see Methods) we had 18 SNPs to use as IVs in the MR analysis. Causal effects estimated by using different meta-analysis methods are reported in Table 4. All of the methods agreed on TMA concentration being causal on *HPS1* expression. To sum up, the estimated causal effect size of *HPS1* on TMA ranged from 0.27 to 0.37 depending on the method, while the causal effect size of TMA on *HPS1* was around −0.09, pointing to the existence of a negative feedback loop.

## Discussions & Conclusion

In this study, we present a metabolome- and transcriptome-wide association study using matching RNA-seq and NMR urine profiles from 555 subjects of the CoLaus cohort. This is the first time such a study is performed on untargeted urine metabolome of healthy individuals. In contrast to targeted approaches that are restricted to a limited set of urine metabolites, our association study uses the binned features of the entire ^1^H NMR spectra as metabolic traits. We identified one gene (*ALMS1*) whose association with two adjacent NMR features around 2.03 ppm is highly significant, surviving even the most conservative correction for multiple hypotheses testing. 16 additional genes are associated with metabolic features with marginal significance with p-values below an adjusted threshold accounting for the estimated number of independent variables (see Table 1). Among the top 17 genes, 10 are in loci with SNPs that have been previously reported as mQTLs. This shows the sensitivity of our study to extract likely candidates of metabolically relevant genes, despite its small sample size and low power.

We used metabomatching to search for promising metabolite candidates underlying gene expression-metabolome features associations. This approach was particularly insightful for our top hit *ALMS1*, as well as the strongest marginally significant association involving *HPS1*: Both genes had previously been implicated by mGWAS linking their loci to compound families. However, in both cases the reported mQTL also harbored other genes, leaving the exact gene-metabolite association ambiguous.

Specifically, the locus associated through mGWAS with N-acetylated compounds includes both *ALMS1* and the *NAT8* gene [18, 27, 36, 39], and the latter seemed to be the more likely candidate due to its known N-acetyltransferase activity. Yet, our association study using transcriptomics data only implicates *ALMS1* and not *NAT8*. Thus, while we cannot rule out a functional role of *NAT8*, the mQTLs of this locus likely act, at least predominantly, as eQTLs through *ALMS1*, pointing to its regulatory role in modulating the compound concentration. This metabolic role of *ALMS1* is also supported through its known role in Alström syndrome characterised by metabolic deficits (PMC6327082) and kidney health disorder phenotypes [40]. Interestingly, in the mGWAS reported by Montoliu et al. using data from a Brazilian cohort, the authors observed the association between N-acetylated compounds and the SNPs located in *ALMS1*/*NAT8* locus with stronger SNP associations in the *ALMS1* gene rather than *NAT8* [39]. They argued that the high ethnic diversity of their study population might have been responsible for breaking down the linkage disequilibrium in the *ALMS1*/*NAT8* region of the genome, resulting in a stronger association for SNPs close or in the *ALMS1* gene compared to other studies.

Our study also sheds more light onto the involved compound: Applying metabomatching on the pseudospectrum from association of all NMR features with the *ALMS1* expression level using a database composed of all N-acetylated compounds NMR spectra, suggested NAA as the best matching metabolite due to the presence of a secondary peak at 7.92 ppm and not missing any high intensity peaks unlike other N-acetylated compounds (Supplementary Figure 5). Interestingly, NAA is the second most abundant metabolite in the brain and involved in neural signalling by serving as a source of acetate for lipid and myelin synthesis in oligodendrocytes [41]. NAA can be detected in urine of both healthy and unhealthy individuals in low concentrations [42] and it has a long history of being a surrogate marker of neural health and a broad measure of cognitive performance [43, 44]. Recently it has been shown that NAA correlates with time measures of neuropsychological performance [45]. The signals of SNPs in *ALMS1* by GWAS with intellectual phenotypes such as self-reported ability in mathematics [46, 47] might therefore be due to its role in modulating NAA. This conjecture of course assumes that NAA levels in relevant brain tissues reflect those in urine and that the *ALMS1* expression variation, and in particular its genetic component, in LCLs or blood, can serve as a proxy for brain tissue. As for *HPS1*, our second strongest association of a gene expression level with urine NMR features, we note that mGWAS previously associated its locus with TMA levels [18, 27, 36]. Yet, most of these studies, including the aforementioned GWAS using a Brazilian cohort [39] considered the *PYROXD2* gene, which is in the same locus, as the most likely modulator of TMA concentrations due to its known function as pyridine nucleotide-disulfide oxidoreductase. While we cannot rule out that this gene is indeed involved in TMA metabolism, in contrast to *HPS1* we have no evidence for association of *PYROXD2* expression levels with TMA. Thus, our data indicates that the mQTLs of this locus act predominantly as eQTLs through *HPS1*, pointing to its regulatory role in modulating TMA.

Our work illustrates the potential of metabolome- and transcriptome-wide association studies for deciphering gene-metabolite relationships. In particular, even with our modest sample size of 555 matched profiles we already had enough power to detect one significant and several marginally significant associations. Moreover, our two strongest associations pinpointed genes in loci implicated by mGWAS as the most likely candidates for transcriptional metabolite regulation. We also showed the possibility of extending correlative work and studying the causal relationship between gene expression levels and metabolite concentrations. Our Mendelian randomization study supported the causal role of *ALMS1* gene expression levels on N-acetylated compound concentration, whereas for *HPS1* we observed a negative feedback loop between its expression levels and TMA metabolite concentrations. Furthermore, this work demonstrated that our metabomatching tool, whose usefulness for elucidating candidate metabolites from mGWAS association profiles [18, 22] as well as auto-correlation signals in NMR data [24] was demonstrated previously, performs equally well on pseudospectra generated by association with gene expression levels.

Our study has many limitations: First, we only had access to gene expression levels of LCLs. While blood and such blood-derived cells are the easiest samples one can obtain from healthy subjects, their expression levels in many cases may only reflect poorly those of the relevant cells and tissues. Furthermore, metabolic reactions are of course driven by enzymes whose protein concentration determines the metabolic rate, and variation in gene expression levels is only one source of variation in active enzyme concentration (next to post-transcriptional and post-translational modifications, as well as their decay rate). Second, metabolite concentrations in urine correspond to excess that is cleared from the body, which depends on food intake and provide a poor proxy for many metabolite concentrations in their relevant location. Nevertheless, our study shows the promise of co-analyzing two or more distinct molecular traits observed in the same cohort.

## Acknowledgements

This work was supported by the Swiss National Science Foundation (grant FN 310030_152724/1) and the NIH (grant R03 CA211815).

## Author Contributions

RS, RR and SB designed the project. RS carried out the computational analysis and prepared the results. BK and RR provided data and feedback on the metabolomics analysis. ZK guided the MR analysis. All authors discussed the results and provided feedback on the manuscript that was written by RS, BK and SB.

