## Supplementary material for "Untargeted metabolome- and transcriptome-wide association study identifies causal genes modulating metabolite concentrations in urine"

### Supplementary Figures

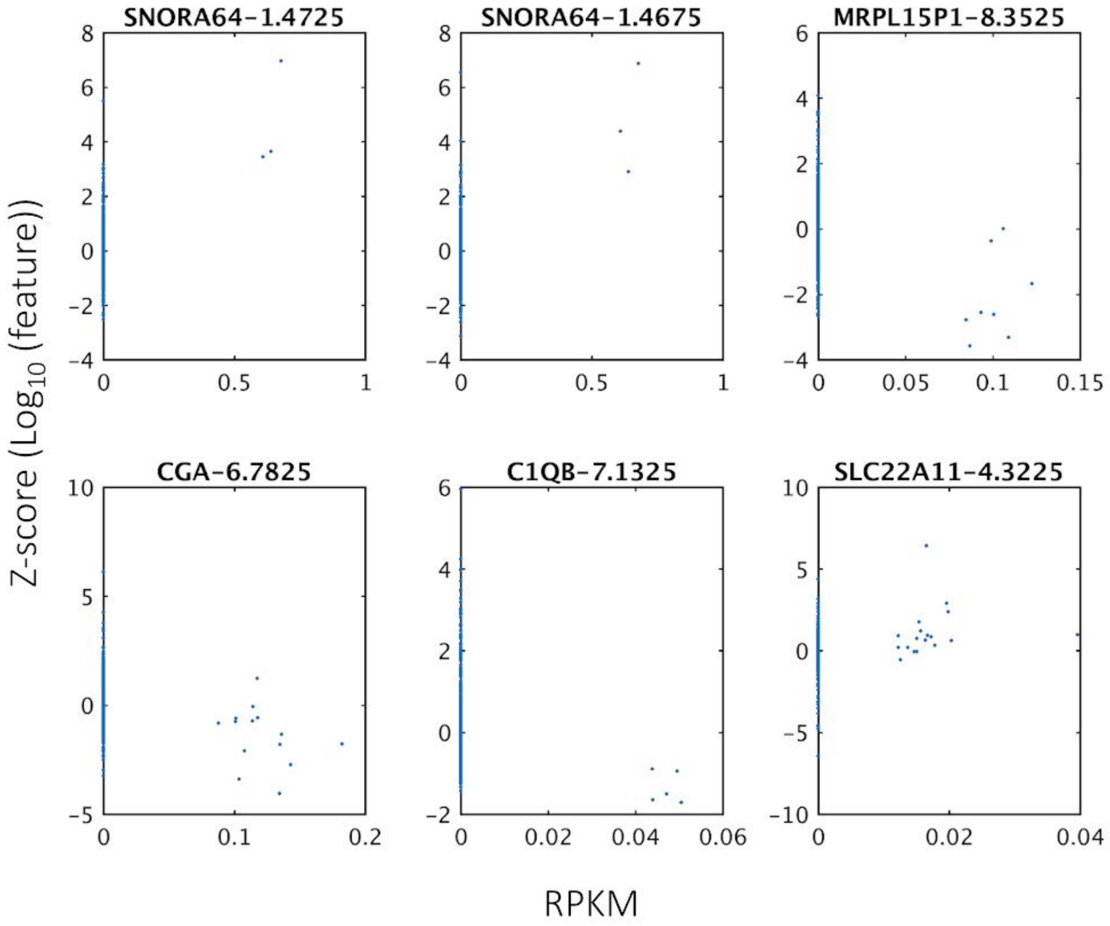


Supplementary Figure 1: Scatter plots of six marginally significant metabolome feature - gene expression associations. X-axis shows the gene's RPKM values and the Y-axis shows the $\log_{10}$ transformed and Z-scored metabolome feature that is associated with the respective gene.

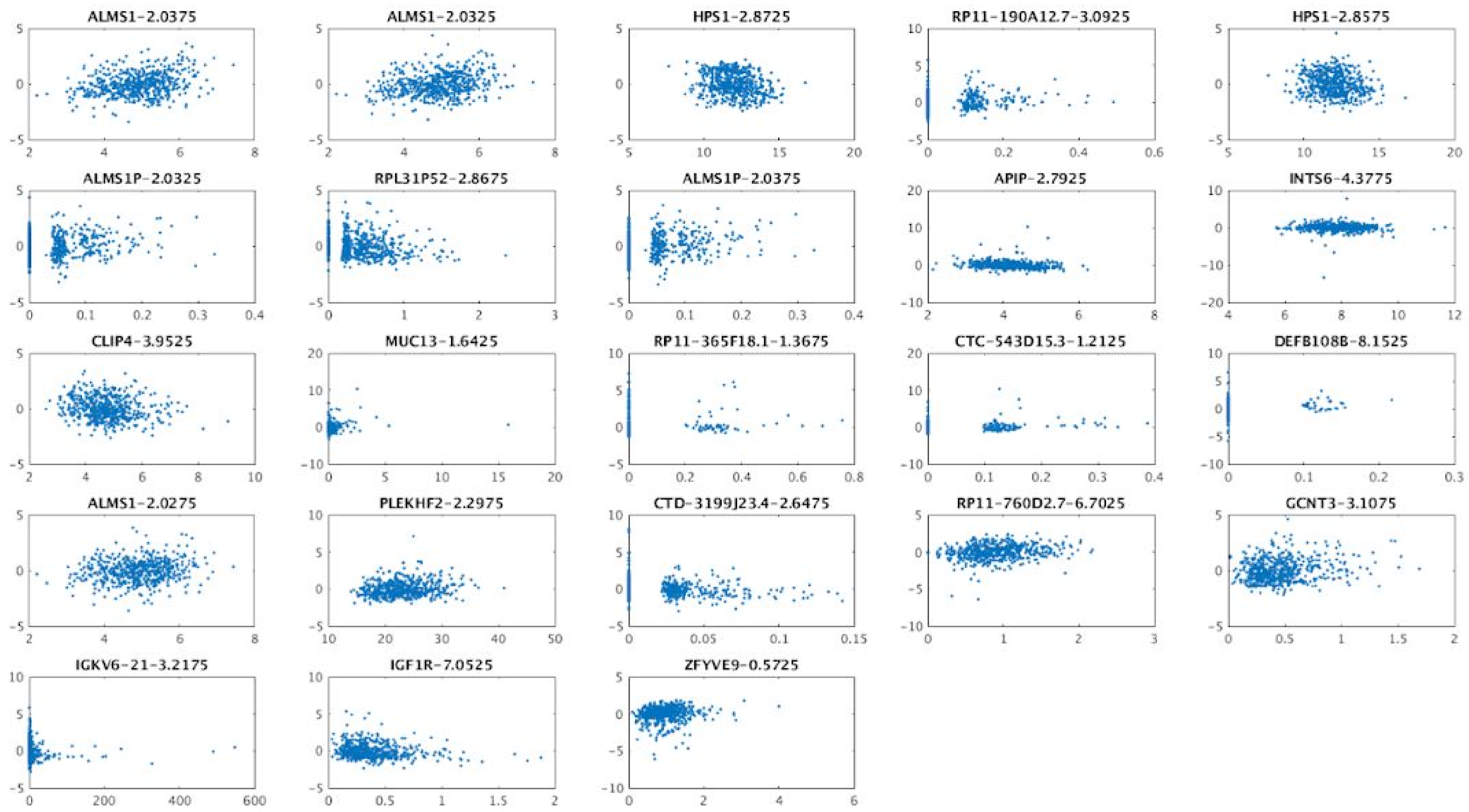


Supplementary Figure 2: Scatter plots of 23 study wide significant metabolome feature - gene expression associations. X-axis shows the gene's RPKM values and the Y-axis shows the $\log_{10}$ transformed and Z-scored metabolome feature that is associated with the respective gene.

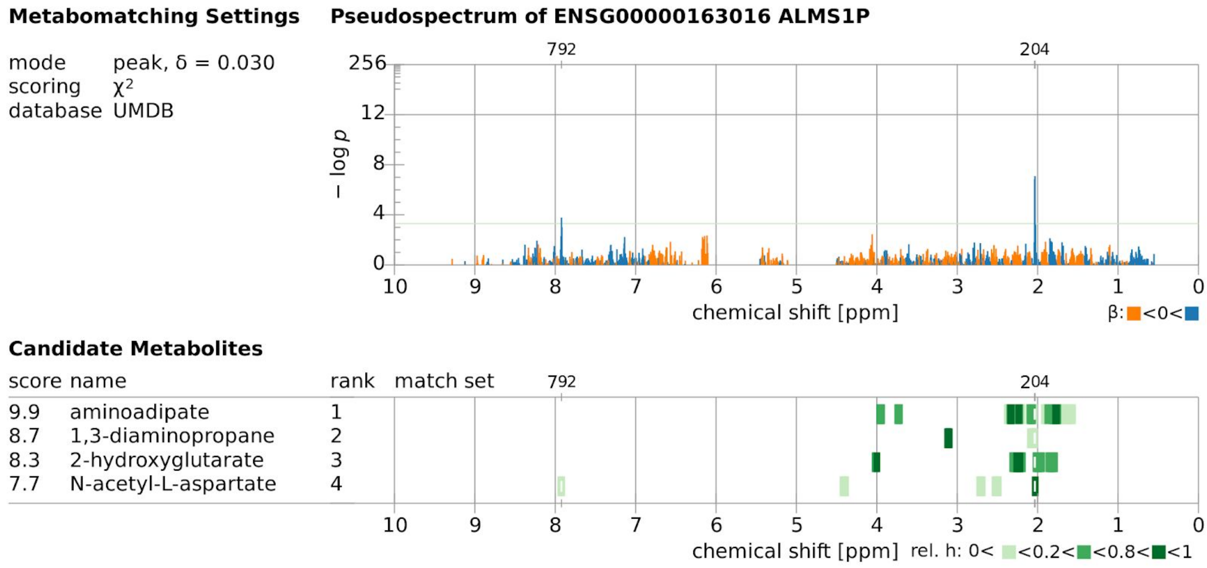


Supplementary Figure 3: CoLaus urine metabolome- ALMS1P gene expression association profile metabomatching figure. Leading features allowing metabolite identification are at 2.0375 ppm, 2.0325 and 7.9225 ppm regions, respectively, which match well with the highest intensity peak of NAA and one of the lower intensity peaks of NAA NMR spectrum, respectively.

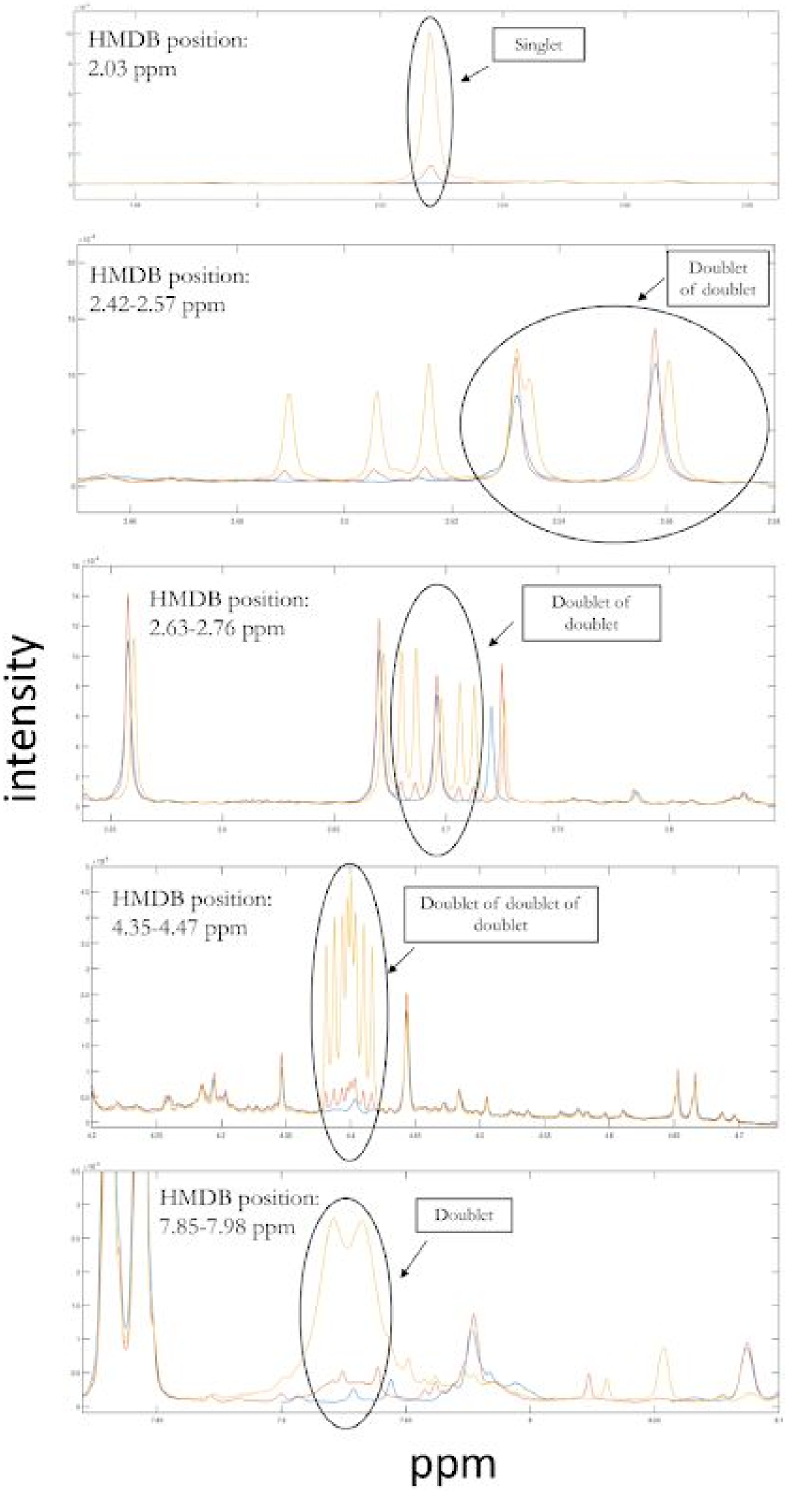


Supplementary Figure 4: Combined NMR profiles of 3 different NMR experiments. Blue spectrum: pooled randomly selected urine samples. Red spectrum: NAA spiked into pooled samples where NAA concentration in the solution is 1 mM. Yellow spectrum: NAA spiked into pooled samples where NAA concentration in the solution is 10 mM.

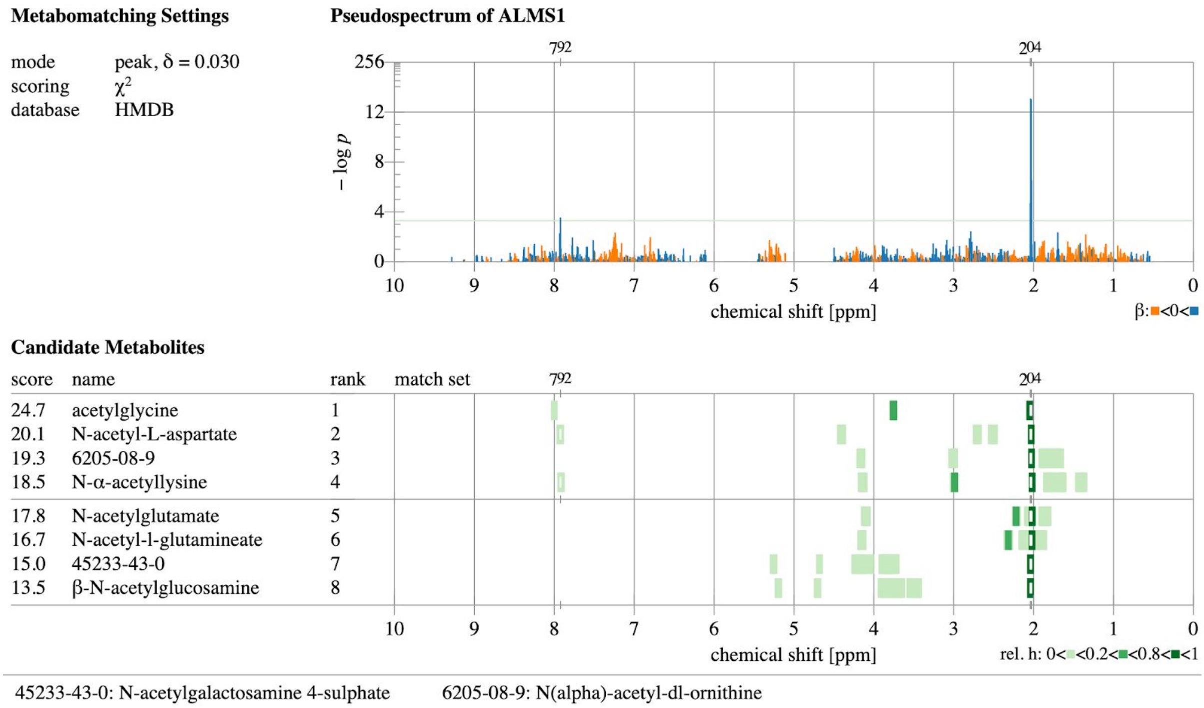


Supplementary Figure 5: ALMS1 metabomatching with all N-acetylated compounds in HMDB and BMRB.

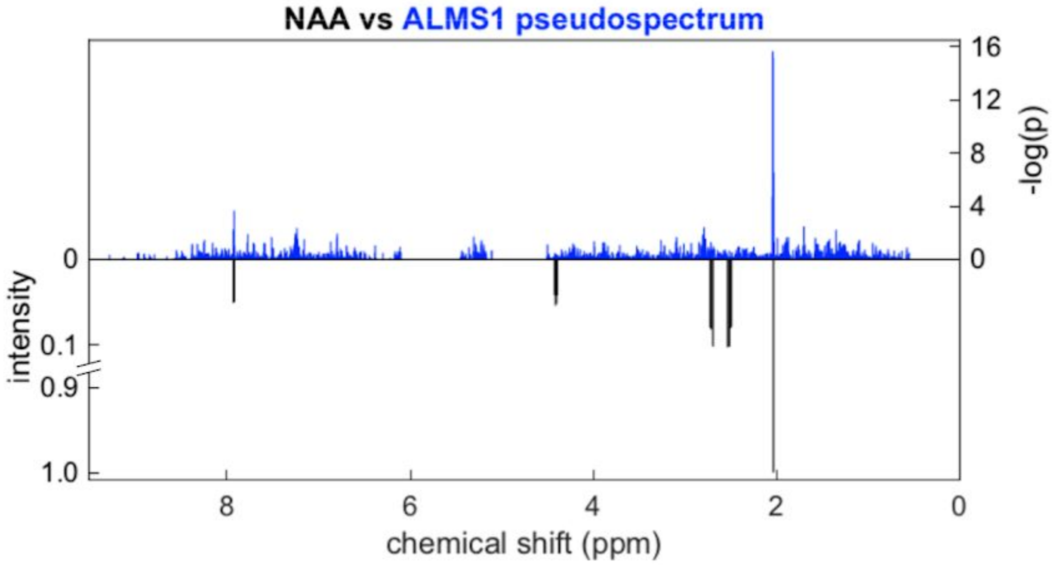


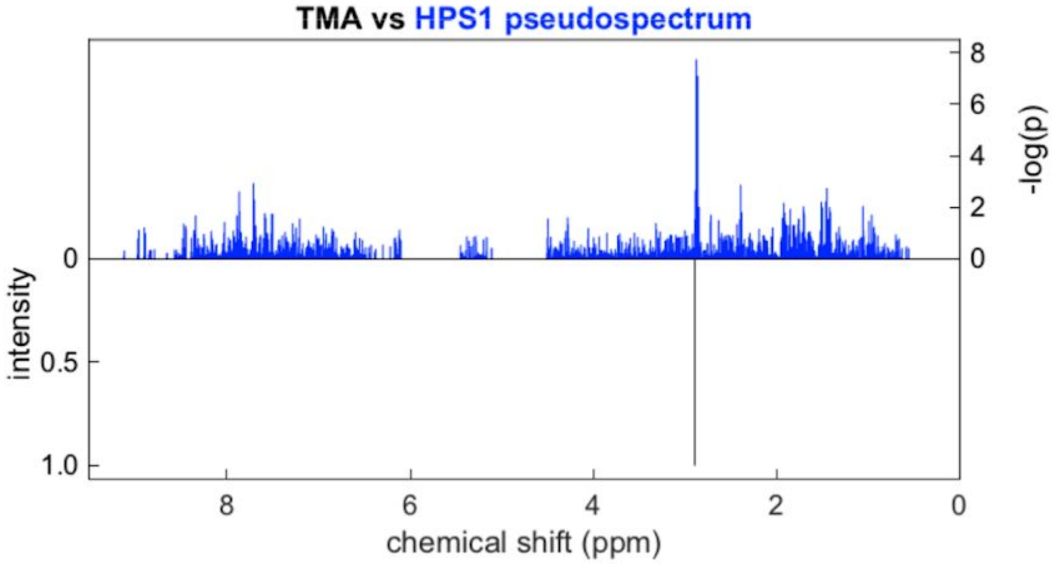


Supplementary Figure 6: Schematic representation of the match between ALMS1 association pseudospectrum and NAA NMR spectrum(top plot) and the match between HPS1 association pseudospectrum and TMA NMR spectrum (bottom plot). Each figure shows -log10 transformed gene expression - metabolome features association p-values on the top and the reference NMR spectrum of the matching metabolite on the bottom. In the NAA - ALMS1 match, leading features allowing metabolite identification are at 2.03 ppm and 7.92 ppm regions which match well with the highest intensity peak of NAA and one of the lower intensity peaks of the NAA NMR spectrum respectively. In the TMA - HPS1 match, leading features allowing metabolite identification are 2.87 and 2.86 ppm which match well with TMA singlet.

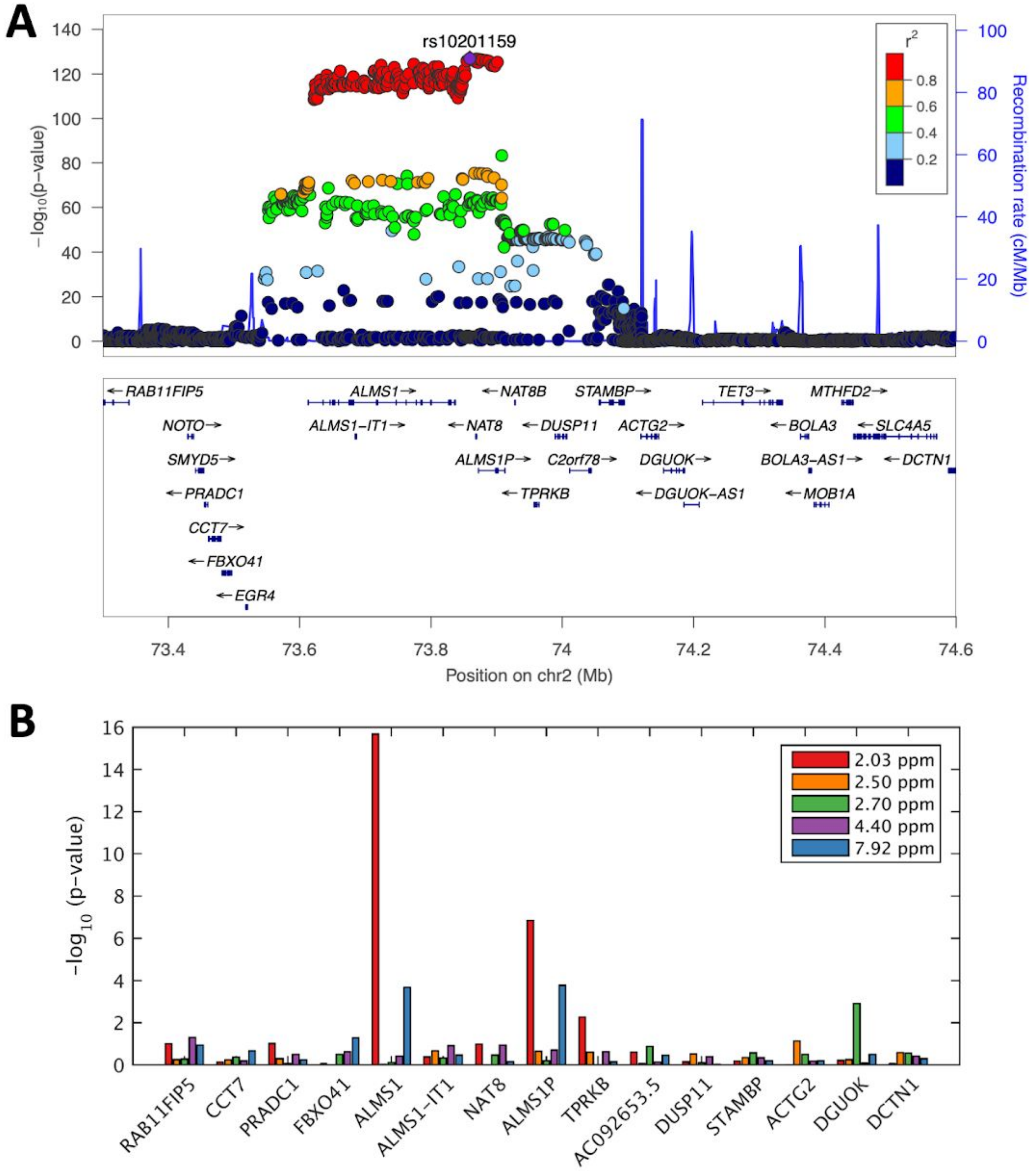


Supplementary Figure 7: A) LocusZoom plot for ALMS1/NAT8 locus, where the SNPs are associated with metabolome feature at 2.0375 ppm, LD colored with respect to lead mQTL. B) Bar plot shows -log10 transformed p-values from associating expression value of 15 genes in the locus with the five NAA features.

### Supplementary Tables

| ALMS1 | shifts | 2.067 | 2.062 | 2.057 | 2.052 | 2.047 | 2.042 | 2.037 | 2.032 | 2.027 | 2.022 | 2.017 | 2.011 |
|---|---|---|---|---|---|---|---|---|---|---|---|---|---|
| *follow-up* | beta | 0.0266 | 0.0625 | 0.0988 | 0.0126 | 0.1019 | 0.2943 | 0.2733 | 0.2094 | 0.0375 | 0.0655 | 0.0126 | 0.0296 |
| | *se* | 0.0588 | 0.0599 | 0.0596 | 0.0582 | 0.0593 | 0.0572 | 0.0579 | 0.0584 | 0.0585 | 0.0580 | 0.0577 | 0.0582 |
| | p-value | 6.51E-01 | 2.97E-01 | 9.83E-02 | 8.29E-01 | 8.71E-02 | 5.12E-07 | 3.73E-06 | 3.92E-04 | 5.21E-01 | 2.60E-01 | 8.27E-01 | 6.12E-01 |
| | r-sq | 0.0704 | 0.0397 | 0.0486 | 0.0874 | 0.0594 | 0.1275 | 0.1087 | 0.0943 | 0.0882 | 0.1030 | 0.1111 | 0.0962 |

| HPS1 | shifts | 2.884 | 2.879 | 2.874 | 2.869 | 2.864 | 2.859 | 2.854 | 2.848 | 2.843 | 2.838 | 2.833 | 2.828 |
|---|---|---|---|---|---|---|---|---|---|---|---|---|---|
| *follow-up* | beta | -0.0285 | -0.0530 | -0.1917 | -0.2492 | -0.1903 | -0.1978 | -0.1290 | -0.0040 | -0.0202 | -0.1004 | 0.0500 | 0.0681 |
| | *se* | 0.0625 | 0.0607 | 0.0606 | 0.0578 | 0.0611 | 0.0610 | 0.0613 | 0.0614 | 0.0610 | 0.0602 | 0.0630 | 0.0618 |
| | p-value | 6.49E-01 | 3.83E-01 | 1.74E-03 | 2.24E-05 | 2.02E-03 | 1.33E-03 | 3.63E-02 | 9.48E-01 | 7.41E-01 | 9.67E-02 | 4.28E-01 | 2.72E-01 |
| | r-sq | 0.0561 | 0.1129 | 0.1174 | 0.1968 | 0.0924 | 0.1028 | 0.0866 | 0.0875 | 0.0989 | 0.1213 | 0.0481 | 0.0825 |

| ALMS1P | shifts | 2.067 | 2.062 | 2.057 | 2.052 | 2.047 | 2.042 | 2.037 | 2.032 | 2.027 | 2.022 | 2.017 | 2.011 |
|---|---|---|---|---|---|---|---|---|---|---|---|---|---|
| *follow-up* | beta | 0.0055 | -0.0274 | -0.0097 | 0.0265 | 0.0225 | 0.1166 | 0.1060 | 0.1237 | 0.0189 | 0.0420 | 0.0331 | 0.0053 |
| | *se* | 0.0643 | 0.0656 | 0.0655 | 0.0636 | 0.0652 | 0.0651 | 0.0655 | 0.0649 | 0.0640 | 0.0635 | 0.0631 | 0.0637 |
| | p-value | 9.31E-01 | 6.76E-01 | 8.82E-01 | 6.78E-01 | 7.30E-01 | 7.45E-02 | 1.07E-01 | 5.75E-02 | 7.67E-01 | 5.09E-01 | 6.00E-01 | 9.34E-01 |
| | r-sq | 0.0697 | 0.0366 | 0.0393 | 0.0879 | 0.0499 | 0.0559 | 0.0467 | 0.0648 | 0.0871 | 0.1003 | 0.1118 | 0.0954 |

Supplementary Table 1: Validation of three essential associations discovered in CoLaus baseline. Association statistics coming from associating CoLaus follow-up urine NMR data with the expression levels of ALMS1, HPS1 and ALMS1P.
